# Genetic insertion of mouse Myxovirus-resistance gene 1 increases innate resistance against both high and low pathogenic avian influenza virus by significantly decreasing replication in chicken DF1 cell line

**DOI:** 10.1101/2024.02.12.579928

**Authors:** Kelsey Briggs, Klaudia Chrzastek, Karen Segovia, Jongsuk Mo, Darrell R. Kapczynski

## Abstract

Avian influenza virus (AIV) is a constant threat to animal health with recent global outbreaks resulting in the death of hundreds of millions of birds with spillover into mammals. Myxovirus-resistance (Mx) proteins are key mediators of the antiviral response that block virus replication. Mouse (Mu) Mx (Mx1) is a strong antiviral protein that interacts with the viral nucleoprotein to inhibit polymerase function. The ability of avian Mx1 to inhibit AIV is unclear. In these studies, Mu Mx1 was stably introduced into chicken DF1 cells to enhance the immune response against AIV. Following infection, titers of AIV were significantly decreased in cells expressing Mu Mx1. In addition, considerably less cytopathic effect (CPE) and matrix protein staining was observed in gene-edited cells expressing Mu Mx1, suggesting Mu Mx1 is broadly effective against multiple AIV subtypes. This work provides foundational studies for use of gene-editing to enhance innate disease resistance against AIV.

## Introduction

Myxovirus-resistance (Mx) proteins are large GTPases that are produced in response to type I interferons (IFN), alpha and beta [1]. Mx proteins are key mediators of the innate antiviral response that directly interact with viral proteins to block replication and/or transcription of several DNA/RNA viruses [2–4]. A functional GTP-binding motif and GTPase activity are necessary for antiviral activity of Mx proteins [5]. Mx proteins self-assemble into oligomeric rings, which are critical for GTPase activity, protein stability, and antiviral activity [6, 7]. Mouse (Mu) Mx1 localizes in the nucleus and is a critical determinant of resistance to influenza A virus (IAV) *in vivo* and *in vitro* [4, 5, 8]. In challenge studies, Mx1-expressing mice are protected from a lethal dose of highly pathogenic avian influenza virus (HPAIV), whereas Mx1-negative mice succumb to infection [9, 10]. Nuclear Mu Mx1 inhibits influenza A viral polymerase activity during primary transcription via interactions with the viral nucleoprotein (NP) [11–13]. The human homologue with antiviral activity, MxA, is found in the cytoplasm, and has a broader range of antiviral activity including IAV, vesicular stomatitis virus (VSV), measles virus, and hepatitis B virus [2, 4, 14].

The role of chicken Mx as an effective antiviral protein is highly contested. Chickens (Ck) have a single Mx gene (Ck Mx) that is induced by type I IFNs [15]. It was originally reported that chicken Mx lacked any antiviral activity *in vitro* [16]. The Ck Mx protein was found to be highly polymorphic with some breeds demonstrating some antiviral activity against IAV and VSV [17, 18]. The Mx residue 631 was implicated in the association with antiviral activity, a serine (S631) lacked antiviral activity, whereas an asparagine (N631) conferred antiviral activity [17, 18]. The susceptible allele (S631) was found in ancestral breeds, but appears to be more prevalent in modern commercial broilers compared to egg-laying strains [19]. Several reports have since negated that S631N conferred antiviral activity *in vitro* and *in vivo* [15, 20–22]. One group reported that Mx chicken alleles containing a N631 did not inhibit IAV replication in chicken embryo fibroblasts (CEFs), and that the cytoplasmic location of Ck Mx was not the reason for the lack of antiviral activity [20, 21]. Schusser et al. reported that Ck Mx lacks GTPase activity, and therefore has no antiviral activity against IAV [15]. An *in vivo* study reported that chickens with N631 had no significant differences in pathology using a low pathogenic avian influenza virus (LPAIV) [23]. However, another *in vivo* study reported that chickens homozygous for N631 had increased resistance early on to a HPAIV, but later succumbed to disease at a higher challenge dose [22]. *In vitro* and *in vivo* experimental results vary on the effectiveness of Ck Mx as an antiviral, but the fact remains chickens are highly susceptible to certain lineages of AIVs.

Poultry vaccination for avian influenza is controversial due to the economic implications, therefore alternate strategies for its control are needed. This study expands upon previous work involving the insertion of Mu Mx1 into the chicken genome. Garber at al., used an avian retrovirus to insert the cDNA encoding the Mu Mx1 protein into CEF cells, which resulted in protection from infection with LPAI viruses [24]. In this work, Mu Mx1 was inserted into an immortalized chicken fibroblast cell line (DF1) using a virus-free transposon vector system to assess resistance to HPAIV and LPAIV replication.

## Methods

### Viruses

All viruses used in this study were propagated in the allotonic cavities of embryonated specific pathogen free (SPF) chicken eggs. Viral titers were determined as previously described [25]. LPAIV included A/Turkey/Wisconsin/1968 H5N9 (Tk/WI), A/Chicken/New Jersey/1997 H9N2 (Ck/NJ), A/Turkey/Virginia/2002 H7N2 (Tk/VA), A/Chicken/California/2003 H6N2 (Ck/CA), and A/Chicken/Pennsylvania/1983 H5N2 (Ck/PA) were utilized in a biosafety level 2 (BSL-2) facility at USNPRC. HPAIV included, A/Turkey/Minnesota/2015 H5N2 (Tk/MN), A/Chicken/Jalisco/2012 H7N3 (Ck/MX), A/American-Wigeon/South Carolina/2021 H5N1 (AW/SC), and A/Turkey/Poland/2021 H5N8 (Tk/PO) and were utilized in a biosafety level 3 (BSL-3) facility at USNPRC.

### Cells

DF1 cells were maintained in flasks containing Dulbecco’s modified eagle medium (DMEM) (ThermoFisher Scientific, Waltham, Ma) with 10% fetal bovine serum (FBS) (Biowest, Bradenton, Fl) and 1% antibiotic-antimycotic (GeminiBio, Sacramento, CA) at 39^°^ C with 5% CO_2_. At 95-100% confluence, cells were passaged using standard procedures.

### Construction of Tol2 plasmid

Sequences were obtained from Genbank for gallus gallus Mx promoter (EF487534) and Mus musculus MX dynamin-like GTPase (NM_01084.1). The Mu Mx1 was *de novo* synthesized in an expression vector (IDT, Newark, NJ). The Mu Mx1 gene and Mx promoter were cloned into the pMAT-miniTol2 vector, containing eGFP and puromycin using Phusion™ (ThermoFisher Scientific, Waltham, Ma) PCR and Gibson Assembly® Cloning Kit (New England Biolabs, Ipswich, MA). The product was transformed in E. coli following the manufacturer’s instructions (Invitrogen, Carlsbad, CA) and plated onto LB agar plates containing carbenicillin (Sigma-Aldrich, Burlington, MA). The plates were grown at 37^°^ C overnight in an incubator. Single colonies were selected and incubated in 5 mL LB broth containing carbenicillin with gentle agitation overnight at 37^°^ C for plasmid isolation. Plasmid DNA was isolated using Quick-DNA™ Miniprep Plus Kit (Zymo Research Irvine, CA) according to the manufacturer’s instructions. Purified plasmid was eluted into 50 µL H_2_O and quantified using the DeNovix DS11-FX spectrometer/fluorometer (DeNovix, Wilmington, DE).

### Construction of DF1 cell line containing Mu Mx1 gene

Two purified plasmids, one containing Mu Mx1 and one containing the Tol2 transposase were transfected into DF1 cells at 50-70% confluency using Xfect transfection reagent (Takara Bio, San Jose, CA) at a 5:1 ratio respectively, according to the manufacturer’s protocol. eGFP was observed at 24 hours post transfection (HPT) using an EVOS M5000 (ThermoFisher Scientific, Waltham, MA). eGFP positive, under puromycin selection, were single cell sorted into a 96-well via fluorescent-activation cell sorting (FACS) at the University of Georgia (Athens, GA), Flow Cytometry Core Center, using a Beckman Coulter Moflo Astrios EQ (Beckman Coulter, Brea, CA). Individual cells were grown in a 96-well until confluency and then expanded into 12-well plates for confirmation studies.

To confirm the presence of the insert in the transfected and sorted cells, cellular RNA was extracted using the RNeasy mini kit (Qiagen, Hilden, Germany) using the manufacturer’s instructions. SuperScript™ III One-Step RT-PCR (ThermoFisher Scientific, Waltham, MA) was used according to the manufacturer’s instruction to amplify Mu Mx1 RNA using Mu Mx1 specific primers. Cells containing the Mu Mx1 plasmid were used as a positive control and plain DF1 cells were used as a negative control.

### Inverse PCR

To determine the location of the insert and confirm it was Tol2 mediated, inverse PCR was used. Splinkerette PCR for mapping transposable elements was adapted for DF1 cells [26]. One microgram of Mu Mx1 expressing DF1 cells was digested with MboI, HaeIII, or AluI in a 20 µl reaction using Cutsmart reaction buffer (New England Biolabs, Ipswich, MA). The digested DNA was self-ligated using T4 DNA ligase (Takara, Japan). The ligated product was precipitated using 3M sodium acetate and 100% ethanol. Phusion PCR (ThermoFisher Scientific, Waltham, Ma) was performed to amplify the 5’ junctions using the following primers 5’f1: 5’-AGT ACT TTT TAC TCC TTA CA-3’ and 5’r1: 5’-GAT TTT TAA TTG TAC TCA AG-3’. A second PCR was performed on the product using the following nested primers 5’f2: 5’-TAC AGT CAA AAA GTA CT-3’ and 5’r2: 5’-AAG TAA AGT AAA AAT CC-3’. The 3’ junctions were amplified using 3’f1: 5’-TTT ACT CAA GTA AGA TTC TAG-3’ and 3’r1: 5’-CTC CAT TAA AAT TGT ACT TGA-3’ followed by a second PCR using nested primers 3’f2: 5’-ACT TGT ACT TTC ACT TGA GTA-3’ and 3’r2: 5’-GCA AGA AAG AAA ACT AGA GA-3’. The inverse PCR products were gel-extracted using the QIAquick Gel Extraction Kit (Qiagen, Hilden, Germany) and sent for sanger sequencing. Sequences were analyzed in Geneious Prime (Geneious, Auckland, New Zealand).

### Virus kinetics

Control DF1 and DF1 expressing Mu1 (DF1xMu) cells were seeded into 12-well plates in quadruplicate. Cells were inoculated with MOI 0.1 of each virus tested at 70% confluency. In brief, the media was removed from the cells and 250 µL of inoculum in plain DMEM was added per well in triplicate. One well was mock infected with plain DMEM. The inoculum was incubated at 39^°^ C with 5% CO_2_ for one hour. After one hour, the inoculum was removed, the cells were washed 2x with phosphate buffered saline (PBS), and 1 mL of fresh DMEM media was added to the cells containing 2% FBS + 1% antibiotic-antimycotic (GeminiBio, Sacramento, CA). Infected cells were incubated at 39^°^C with 5% CO_2_ and supernatants (0.1 mL) were taken at either 2-, 24-, and 48-hours post infection (HPI) or at 3-, 5-, 10-, 24-, 48-, and 72-HPI for detection of virus by Quantitative real-time RT-PCR (qRT-PCR). Studies involving LPAIV include trypsin in the medium at a concentration of 1-2 µg/mL as previously described [27]. For the early induction of Mx experiments, Ck IFNα (Bio-Rad, Hercules, CA) was added to the media 18 hours prior to infection at 1000 units/mL. Cells were imaged at 24- and 48-HPI using an EVOS M5000 (ThermoFisher Scientific, Waltham, MA) to observe cytopathic effect (CPE). Each experiment was performed three independent times.

RNA extractions were performed using Ambion Magmax kit (ThermoFisher Scientific, Waltham, MA) on a KingFisher Flex system (ThermoFisher Scientific, Waltham, MA) following the manufacturer’s instructions. RT-qPCR was used to detect and determine virus titers using the AgPath ID one-step RT-PCR kit (ThermoFisher Scientific, Waltham, MA) from the cell culture supernatants as previously described [28]. A standard curve of RNA from titrated AIV virus stock was run to establish titer equivalents of virus. Values were normalized to the DF1 controls cells. Percent reduction was calculated dividing individual titers of virus grown in the DF1xMu cells by the average control values. The product was then multiplied by 100 to get a percent reduction compared to the DF1 control.

### Immunofluorescence

Cells were seeded into an I-Bidi 8-well chambered slide (Fisher Scientific, Waltham, MA). At 60-70% confluency, cells were infected with MOI 0.1 of virus as described above. At 24 HPI, the media was removed, and the cells were fixed for 5 minutes with 1:1 ice cold ethanol:methanol. After fixation, cells were then washed 2x with PBS and blocked with 3% bovine serum albumin (BSA) for 1 hour at room temperature (RT). Following two washes with 1% BSA, the primary antibody, mouse monoclonal anti-matrix antibody (Southern Biotech, Birmingham, AL), was diluted 1:500 and added to the cells in 1% BSA for 1 hour at RT. Cells were washed 3x with 1% BSA and incubated with the secondary antibody, goat-anti-mouse IgG H&L Cy3 (Abcam, Cambridge, United Kingdom) diluted to 1:500 in 1% BSA for 1 hour at RT. Cells were washed 3x with 1% BSA and counterstained with NucBlue Ready Probes (DAPI) (Invitrogen, Carlsbad, CA) for 10 minutes. Immunofluorescence was visualized with an EVOS M5000 (Invitrogen, Carlsbad, CA).

### Mx Modeling

Predicted structures for Mu Mx1 and Ck Mx were created using Distance-Guided-Protein Structure Prediction (D-I-TASSER) an online prediction software created by the Zhang lab at University of Michigan using the following link [29]. https://zhanggroup.org/D-I-TASSER/. The structures were visualized and analyzed using ChimeraX version 1.6.1 (UCSF, San Fransisco, CA) [30].

### Alignments

All alignments were performed in Geneious Prime using MUSCLE 5.1 alignment (Geneious, Auckland, New Zealand). The consensus sequence is shown at the top and a dot represents a conserved residue. Mx proteins were obtained from Genbank: Hu MxA (NP_001171517.1), Mu Mx1 (NP_034976.1), Ck S631 (NP_989940.2), and Ck N631 (ABH07790.1). Virus sequences were obtained from GISAID [31].

### Statistical analysis

Viral titers were compared with either an unpaired T-test or two-way ANOVA with Tukey multiple comparison (GraphPad Prism, San Diego, CA). Different lower-case letters indicate statistical significance between compared groups. All statistical tests used p < 0.05 as being statistically significant.

## Results

### Development of chicken cell line expressing Mu Mx1

These studies were designed to stably insert the Mu Mx1 gene, under control of the virus-inducible chicken Mx promoter, into the chicken DF1 cell genome to enhance disease resistance to avian influenza via the innate immune response. The Ck Mx and Mu Mx1 proteins contain 705 and 631 amino acids (AA) respectively, that are approximately 41.9% identical. Interestingly, there are 412 AA differences between them, however the GTP-binding consensus motifs contain identical residues, which suggests Mu Mx1 GTPase activity would be functional in DF1 cells (Supplemental Figure 1 and Supplemental Table 1). A Tol2 transposon was used to deliver the Mu Mx1 gene using a plasmid-based system. The insert also contained eGFP selection marker under a CMV promoter for downstream sorting and purification (Figure 1). The constructs were single cell sorted via FACS by eGFP and grown to confluency, additional purifications were performed as needed (Figure 2A). Single cell sorting and grow out produced single Mu Mx1-expressing clones. Detection of the insert was confirmed by RT-PCR using primers specific for Mu Mx1 (Figure 2B). The insert was mapped to chromosome 3 in DF1 cells using inverse PCR (Figure 2C).

**Figure 1.**
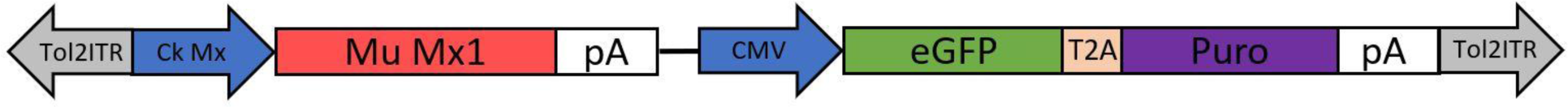
Schematic of insertion in pMAT-minitol2 plasmid. The Tol2ITR are inverted terminal repeat sequences, the sequences within the repeats were inserted into the DF1 genome. The Ck Mx promoter sequence and Mu Mx1 gene were obtained from Genbank, *de novo* synthesized, and cloned into the pMAT-minitol2 vector. eGFP linked to puromycin was added for easy detection and downstream selection.

**Figure 2.**
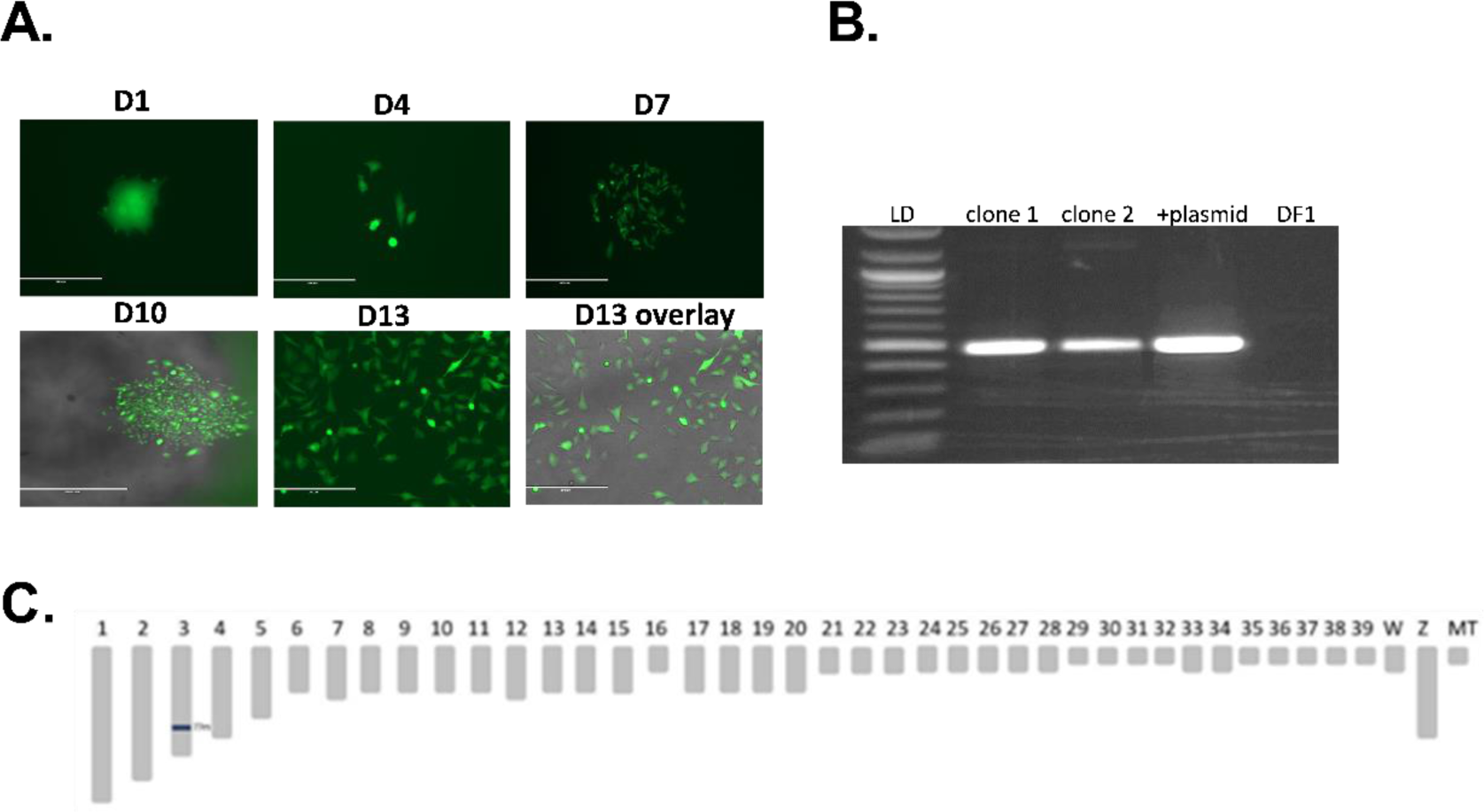
Detection and location of Mu Mx1 insertion in DF1 cells. (A) DF1 cells containing Mu Mx1 were single cell sorted via eGFP under puromycin selection and monitored until confluency. At day 13, an overlay of eGFP and transmitted light was performed using a fluorescent microscope to show a homologous population. (B) RNA was extracted from two confluent clones to confirm expression of Mu Mx1 in the DF1 cells. Transfected plasmid was used as a positive control and DF1 cells were used as a negative control. (C) The location is the Tol2 insert was determined using inverse PCR. The schematic represents a simplified chicken karyotype. The insert was found to be in chromosome 3 at position 77394060.

### Low pathogenic avian influenza virus titers are decreased in cells expressing Mu Mx1

DF1xMu and normal control DF1 cells were infected with four subtypes of LPAI in the presence of trypsin. In the DF1xMu cells, Tk/WI had the most significant reduction in replication with viral titers reduced by 95% compared to the DF1 control cells at 24 HPI (Figure 3A). The viral titers of Ck/NJ were reduced by 90% when grown in the DF1xMu cells compared to the DF1 control cells (Figure 3B). Both Tk/VA and Ck/CA viral titers demonstrated greater variability in reductions with titers reduced by approximately 55% and 70% respectively, in the DF1xMu cells (Figure 3C and 3D). Despite the variation in reduction, all virus isolates were significantly reduced in the DF1xMu cells, confirming that Mu Mx1 is an effective anti-viral protein against AIV.

**Figure 3.**
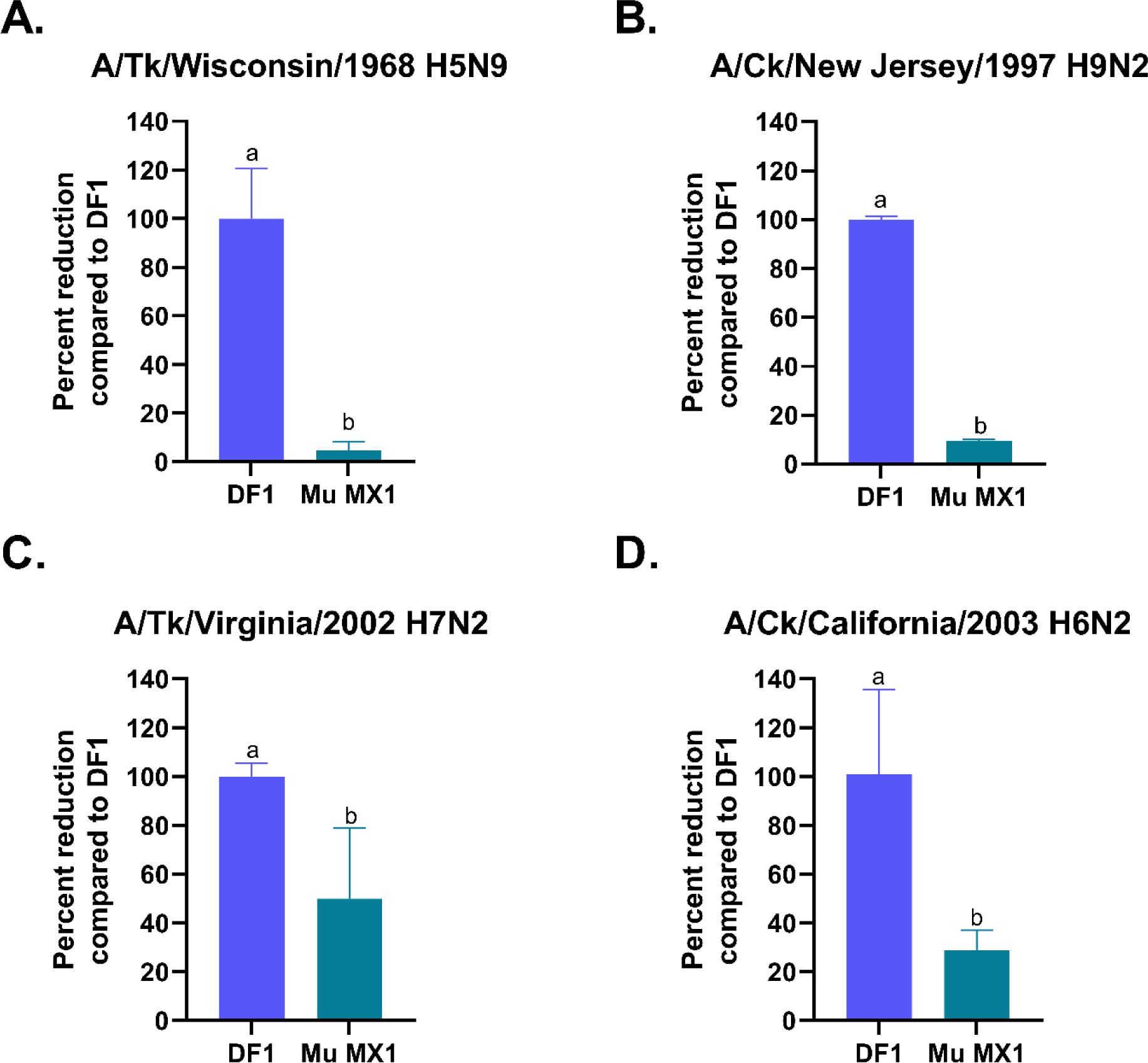
Growth of LPAI viruses in DF1 cells expressing Mu Mx1 and DF1 control cells. DF1xMu and DF1 cells were infected with a LPAI virus at MOI 0.1. Supernatants were taken at 24 HPI for RNA extraction and viral titers were determined by RT-qPCR. The standard deviation of three independent replicates is shown for each cell line and virus. An unpaired T-test was performed to determine statistical significance. p<0.05 is considered statistically significant and is denoted by a different letter (a,b). Values were normalized to the titers of the DF1 control cells. Titers grown in the DF1xMu cells are shown as the percent reduction compared to the control. A representative graph is shown for each virus in triplicate, three independent experiments were performed. Four different LPAI viruses were used (A) A/Turkey/Wisconsin/1968 H5N9 (Tk/WI), (B) A/Chicken/New Jersey/1997 H9N2 (Ck/NJ), (C) A/Turkey/Virginia/2002 H7N2 (Tk/VA), and (D) A/Chicken/California/2003 H6N2 (Ck CA).

### Highly pathogenic avian influenza virus titers are decreased in cells expressing Mu Mx1

Growth curves with HPAIV were also performed in the DF1xMu cells to determine effectiveness against two HPAI isolates. Tk/MN replicated at significantly lower levels in the cells expressing Mu Mx1. At 2 HPI, viral titers were comparable, however at 24 and 48 HPI titers were over 1log_10_ lower (90% reduction) in the DF1xMu cells (Figure 4A). At 24 HPI, Ck/MX viral titers were reduced by almost 4log_10_ (99.99% reduction) in the DF1xMu cells. At 48 HPI, titers increased, but were still 2log_10_ lower than in the control cells (Figure 4B). CPE for each virus and cell line was observed at 24 HPI. HPAIV viruses had substantially less CPE in the DF1xMu cells (Figure 4C). Immunofluorescence performed at 24 HPI also demonstrated less matrix protein staining in the DF1xMu cells compared to the DF1 control cells (Figure 4D).

**Figure 4.**
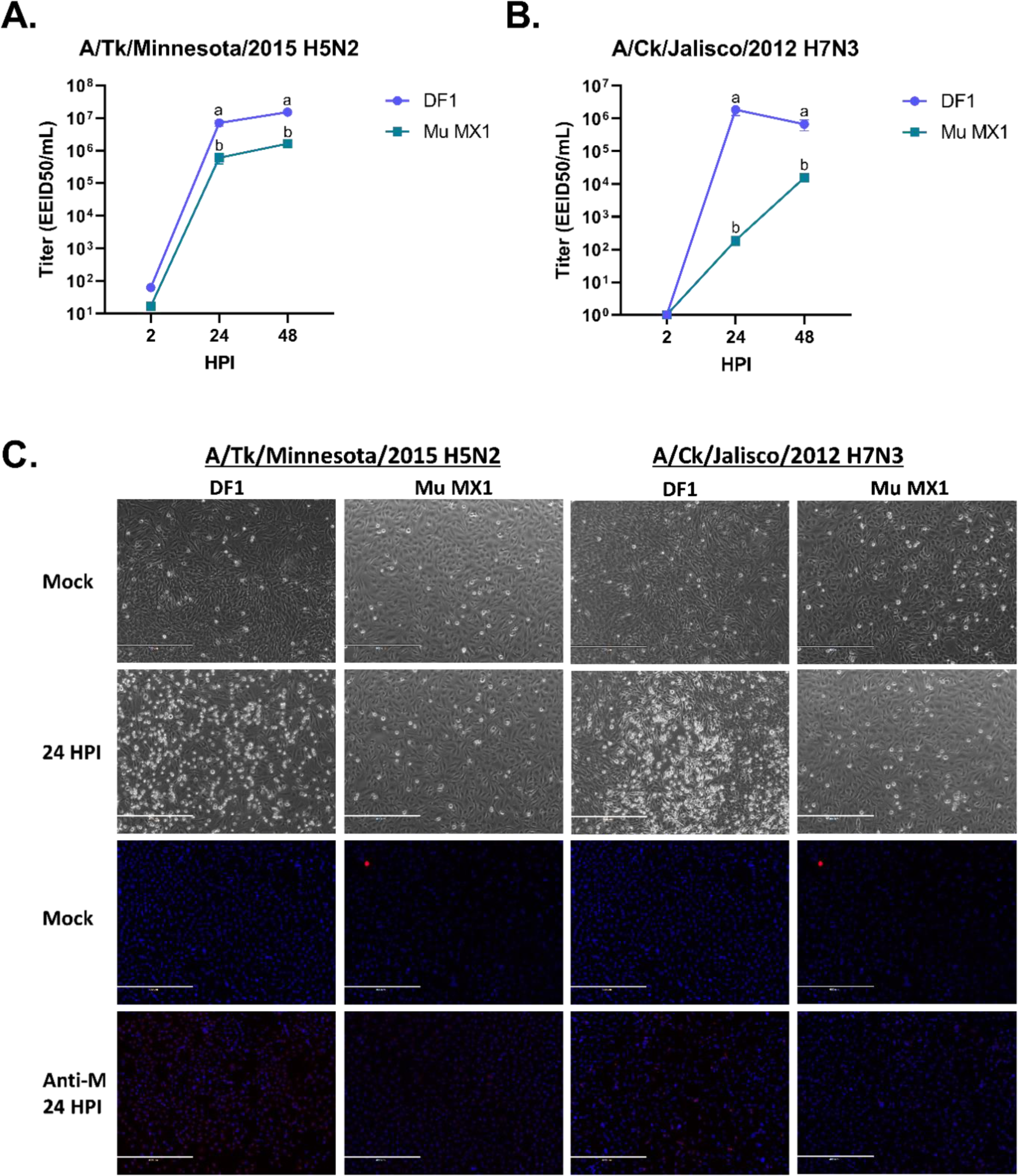
Growth of HPAI viruses in DF1 cells expressing Mu Mx1 and DF1 control cells. (A,B) DF1xMu and DF1 cells were infected with a HPAI virus at MOI 0.1. Supernatants were taken at 2-, 24-, and 48-HPI for RNA extraction and viral titers were determined by RT-qPCR. The standard deviation of three independent replicates is shown for each cell line and virus. Tukey’s multiple comparison test was performed to determine statistical significance at 24- and 48-HPI infection. p<0.05 is considered statistically significant and is denoted by a different letter (a,b). A representative graph is shown for each virus in triplicate, three independent experiments were performed. (A) A/Turkey/Minnesota/2015 H5N2 (Tk/MN), a 2.3.4.4c isolate, and (B) A/Chicken/Jalisco/2012 H7N3 (Ck/MX), a Mexican isolate were used. (C) Cells were inoculated with MOI 0.1 of each virus in glass chamber slides. At 24 HPI, monolayers were observed for CPE and virus was detected using a mouse monoclonal anti-matrix antibody. M protein staining was observed using a goat-anti-mouse Cy3 secondary antibody and nuclei were counterstained with DAPI. Immunofluorescence was visualized with an EVOS M5000 microscope. A representative mock infection picture was used for comparison.

Two recent 2.3.4.4b HPAIV isolates were also tested in DF1xMu and control cells to measure the percent reduction of virus at 24 HPI. Virus replication of the AW/SC isolate was reduced by 80%, whereas the Tk/PO isolate was reduced by 68% compared to virus grown in the control cells (Figure 5A and 5B).There was a noticeable difference in the amount of CPE observed in the control cells and the DF1xMu cells. Both 2.3.4.4b viruses had decreased amounts of CPE at 24 HPI in the DF1xMu cells compared to the control cells. At 48 HPI, the viruses grown in the DF1xMu cells maintained a monolayer, albeit with observed CPE, whereas most of the control cells were destroyed (Figure 5C). Similar to the CPE results, the DF1xMu cells had more visible nuclei but less matrix protein staining than the control cells (Figure 5D). Overall, HPAI viruses grown in cells containing Mu Mx1 had significantly decreased viral titers, less CPE, and lower amounts of matrix protein present.

**Figure 5.**
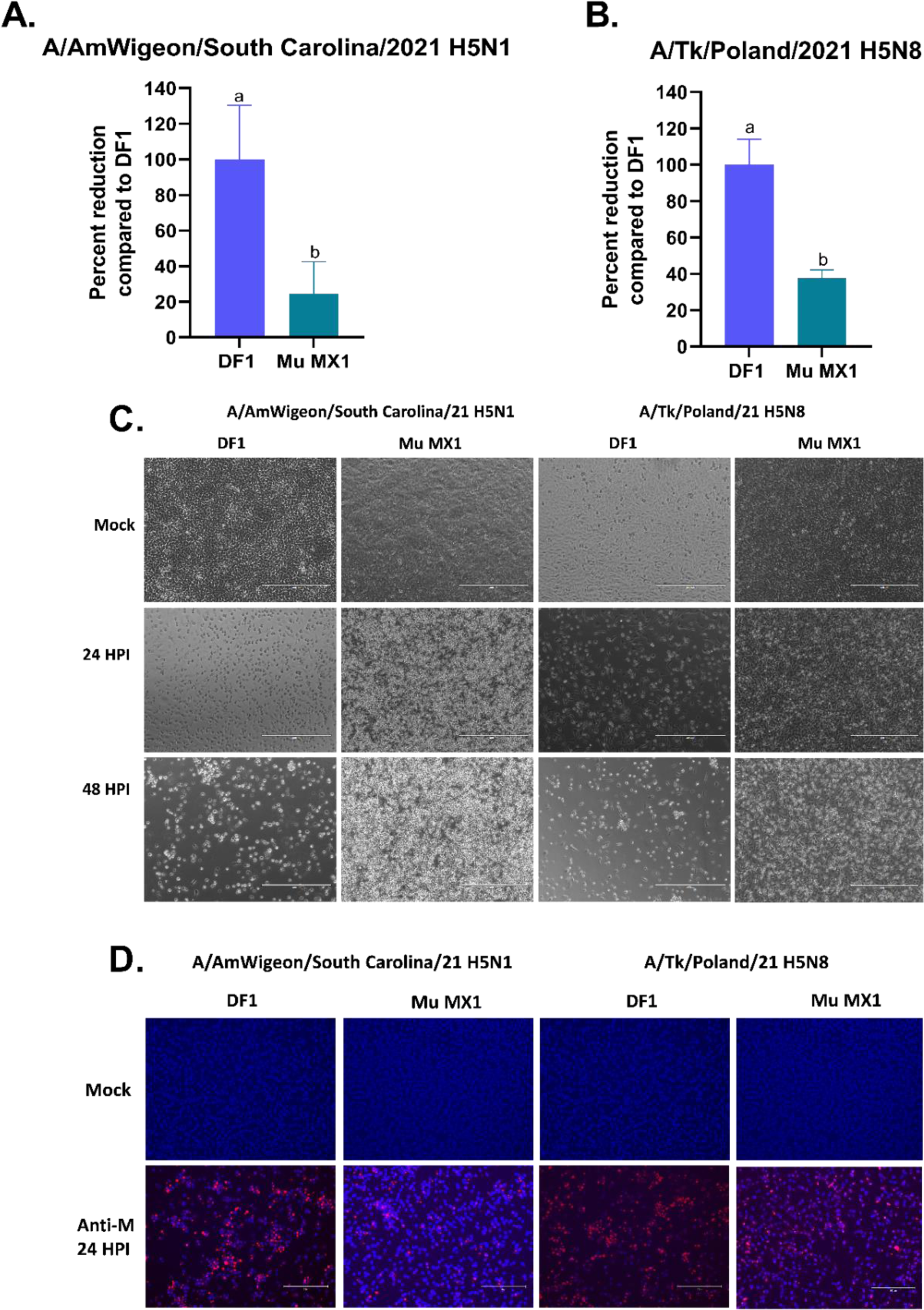
Growth of 2.3.4.4b HPAI viruses in DF1 cells expressing Mu Mx1 and DF1 control cells. DF1xMu and DF1 cells were infected with a 2.3.4.4b isolate, (A) A/American Wigeon/South Carolina/2021 H5N1 (AW/SC) and (B) A/Turkey/Poland/2021 H5N8 (Tk/PO) at MOI 0.1. Supernatants were taken at 24 HPI for RNA extraction and viral titers were determined by RT-qPCR. The standard deviation of three independent replicates is shown for each cell line and virus. An unpaired T-test was performed to determine statistical significance. p<0.05 is considered statistically significant and is denoted by a different letter (a,b). Values were normalized to the titers of the DF1 control cells. Titers grown in the DF1xMu cells are shown as the percent reduction compared to the control. A representative graph is shown for each virus in triplicate, three independent experiments were performed. (C) CPE was observed in cell lines at 24- and 48-HPI for both isolates used. (D) Cells were inoculated with MOI 0.1 of each virus in glass chamber slides. At 24 HPI, virus was detected using a mouse monoclonal anti-matrix antibody. M protein staining was observed using a goat-anti-mouse Cy3 secondary antibody and nuclei were counterstained with DAPI. Immunofluorescence was visualized with an EVOS M5000 microscope. A representative mock infection picture was used for comparison.

### Pretreatment with Ck IFNα decreases viral titers in cells expressing Mu MX1

To determine if prior activation enhanced innate immunity against AIV, we preincubated Ck IFNα with DF1xMu and control cells 18 hours prior to infection. Results demonstrate Ck/CA and Ck/PA viral titers were significantly reduced at 24 and 48 HPI when pretreated with IFNα in both DF1xMu cells and DF1 cells (Figure 6). LPAI viral titers were the most reduced in the DF1xMu + IFNα. Interestingly no statistical difference in titer was observed between Df1xMu and control DF1s pretreated with IFNα (Figure 6).

**Figure 6.**
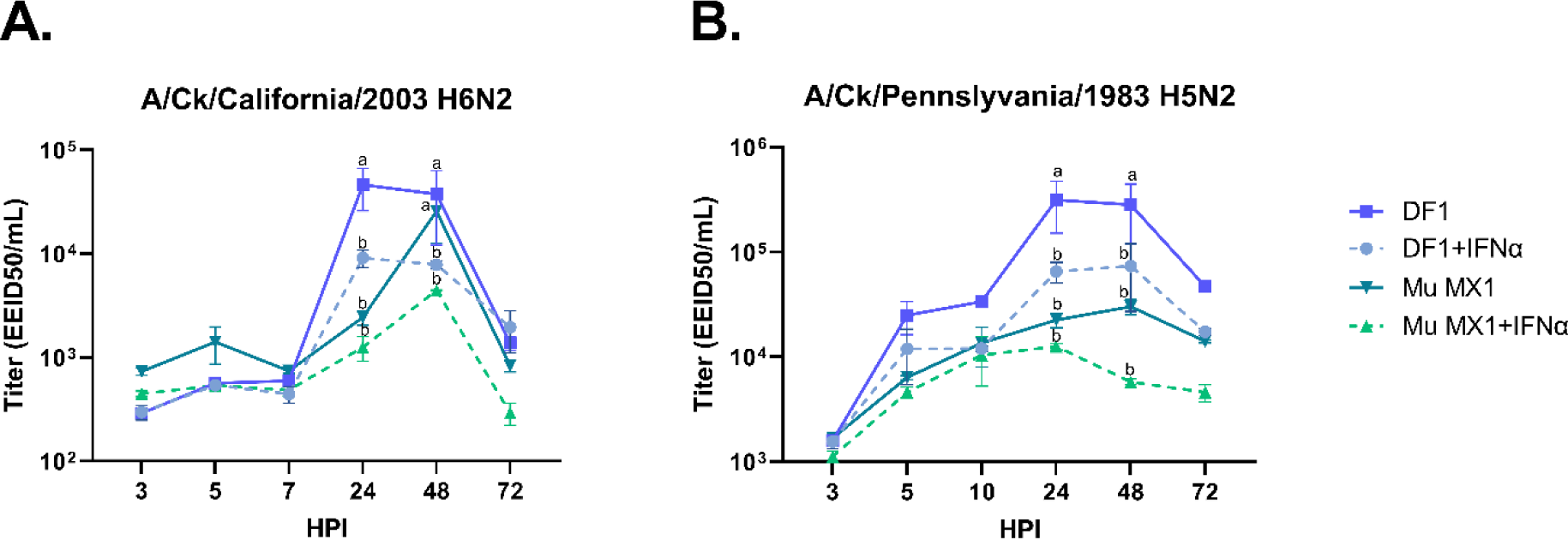
Growth of LPAI viruses in Mu Mx1 and DF1 cells in the presence of Ck IFNα. DF1xMu and DF1 cells were pretreated with Ck INFα 18 hours prior to infection. Monolayers were infected with a LPAI virus at MOI 0.1. Supernatants were taken at 3-, 5-, 7-, 24-, 48-, and 72-HPI for RNA extraction and viral titers were determined by RT-qPCR. The standard deviation of three independent replicates is shown for each cell line and virus. Tukey’s multiple comparison test was performed to determine statistical significance at 24-, 48-, and 72-HPI infection. p<0.05 is considered statistically significant and is denoted by a different letter (a,b). A representative graph is shown for each virus in triplicate, three independent experiments were performed. (A) A/Chicken/California/2003 H6N2 (Ck/CA) and (B) A/Chicken/Pennsylvania/1983 H5N2 (Ck/PA) were used.

### Structural differences between viral and host Mx proteins

Because Mx interacts with viral NP proteins, we next examined the structural differences between viral NP, PB2, PB1, and PA protein sequences from the isolates used here (Table 1). The older Tk/MN and Ck/MX isolates were compared against the two recent 2.3.4.4b isolates, AW/SC and Tk/PO. All proteins examined were approximately 99% similar, however some notable differences were observed. Two differences were observed in NP, I/M105V and N450S and no differences were observed in PB2 (Table 1). PB1 contained three differences, S59T, S375V, and N694S, while PA contained one difference, P400S (Table 1). These differences may impact the susceptibility to Mu Mx1.

**Table 1.** Differences in replication machinery amino acid sequences between current 2.3.4.4b HPAI viruses and contemporary HPAI viruses used in study.

| NP | PB2 | PB1 | PA |
| --- | --- | --- | --- |
| I<sup>1</sup>/M<sup>2</sup>105V |  | S59T | P400S |
| N450S |  | S375V |  |
|  |  | N694S |  |

1-A/Tk/Minnesota/15 H5N2 2-A/Ck/Jalisco/12 H7N3

The D-I-TASSER protein structure prediction server was used to predict the structures of Mu Mx1 and Ck Mx, and ChimeraX was used to visualize and superimpose the proteins via alignment (Figure 7) [29, 30]. Ck Mx contains approximately 80 amino acids in the N-terminal not found in mammalian Mx proteins so an accurate structure could not be predicted in that region (Figure 7). The region with the greatest structural homology, shown in black, was the GTPase domain, at amino acids 90-378 (Figure 7 and Supplemental Figure 2). There was a small portion in the stalk domain that was also conserved in the C-terminal from amino acids 644-651 (Figure 7 and Supplemental Figure 2). Influenza antiviral activity has been attributed to the L4 loop (grey box) in Mu Mx1, while the regions do not share structural homology, the Ck Mx 631 residue is located just downstream (Figure 7 and Supplemental Figure 2) [32]. Although Mu Mx1 and Ck Mx appeared to be structurally similar, there were notable differences that could impact the interaction of the protein with viral NP.

**Figure 7.**
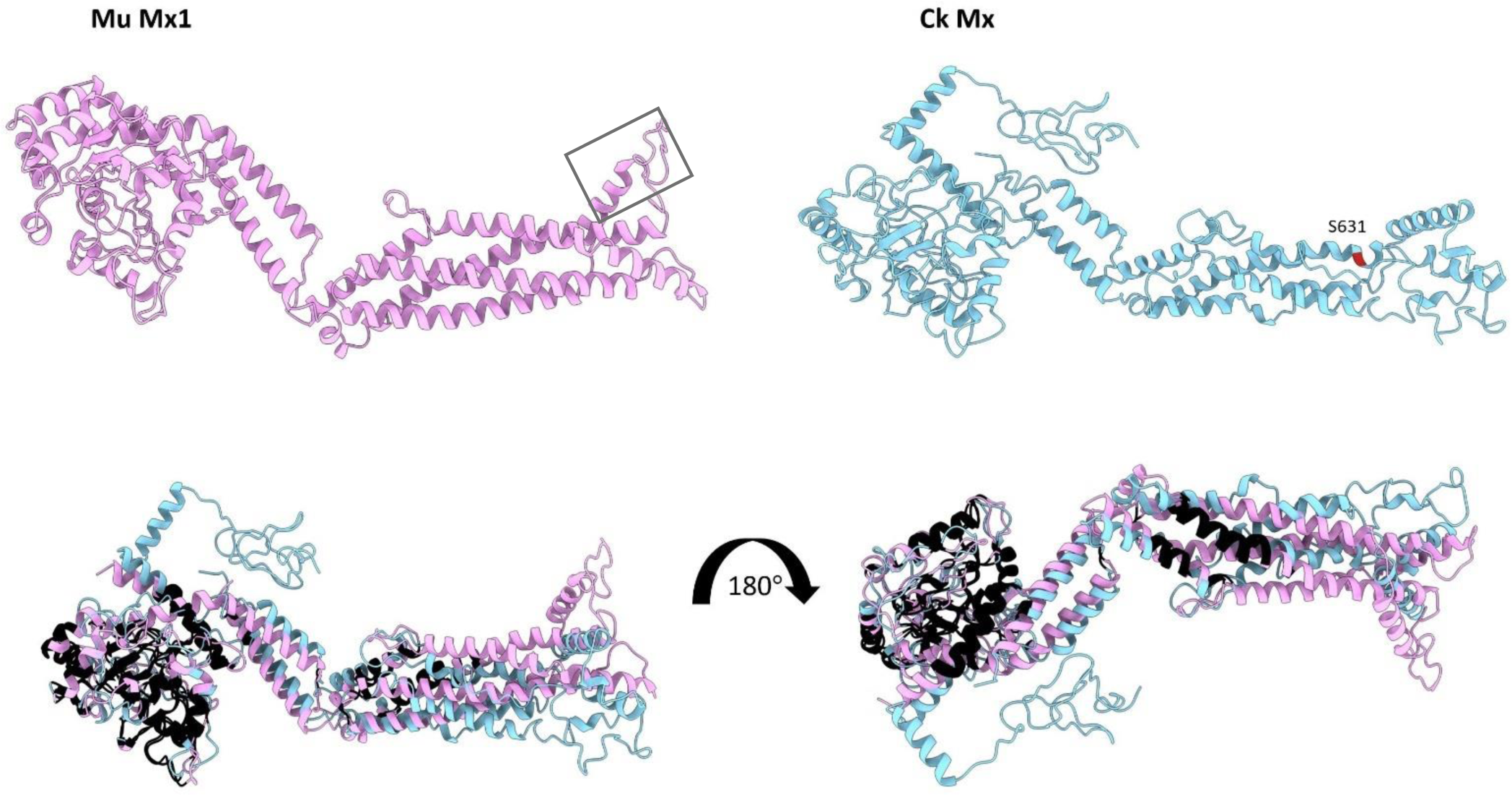
Structural differences between Mu Mx1 and Ck Mx. Predicted protein structures were developed by the D-I-TASSER prediction server. The Mu Mx1 protein structure is in pink and the Ck Mx is in light blue. Residue S631 is labeled in red in Ck Mx. The grey box denotes the L4 loop in Mu Mx1. ChimeraX was used to visualize and overlay the proteins. Regions of homology are colored black.

## Discussion

The goal of this work was to create an immortalized DF1 cell line containing a functional Mu Mx1 that could enhance disease resistance to avian influenza. A miniTol2 transposon system was used to insert Mu Mx1 into DF1 cells and expression was confirmed via RT-PCR and microscopy. Expression of Mu Mx1 in chicken DF1 cells was able to significantly decrease viral growth of LPAI and HPAI viruses. Several AIV subtypes were tested in this study (H5, H6, H7, and H9) suggesting Mu Mx1 can broadly enhance the innate immune response to AIV. Additionally, the presence of Ck IFNα enhanced Mu Mx1’s ability to inhibit viral growth. Structural analysis identified few regions of homology between Mu Mx1 and Ck Mx, however the GTP-binding consensus motifs were conserved. Taken together, this work demonstrates that chicken innate immunity can be improved upon using Mu Mx1 to help combat AIV infections.

Mx1 proteins inhibit IAV replication by interacting with the PB2 and NP proteins [12, 33–36]. It is thought that the Mx1 oligomeric ring binds the IAV ribonucleoprotein complex disrupting the PB2-NP interaction by a GTPase-mediated mechanism [32, 36]. The origin of the NP protein plays a pivotal role in Mx1 sensitivity. Human and mouse IAV strains tend to be more resistant, whereas avian strains are typically more susceptible to Mu Mx1 [12]. All viruses used in this study were of avian origin, yet there were notable differences in Mu Mx1 susceptibility between the HPAI viruses used. There was a considerable difference in the amount of CPE observed between the 2.3.4.4b viruses and the older HPAI isolates. Both 2.3.4.4b viruses caused substantial CPE at 24 HPI and caused a complete removal of the monolayer at 48 HPI in control cells, whereas the older strains caused less CPE. The lack of a monolayer in the control cells infected with 2.3.4.4b viruses likely negatively affected the viral titers, which resulted in a less significant decrease when compared to the DF1xMu cells.

Another explanation for the difference in HPAI susceptibility could be from sequence variation in the replication machinery. There were 6 changes between the 2.3.4.4b viruses and the older isolates. A valine (V) at position 105 of NP has been shown to be responsible for increased pathogenicity in chickens and is proposed to be an adaption from wild birds to poultry [37]. Both the 2.3.4.4b viruses, AW/SC and Tk/PO, tested in this study contained a V at position 105 in NP, whereas Tk/MN and Ck/MX did not. Riegger et al. found that a cluster of amino acids in NP (52, 100, 283, and 313) were responsible for Mu Mx1 and Hu MxA sensitivity, position 105 is located near that cluster and may also play a role in Mx sensitivity (data not shown) [38]. Another difference was at position 450 in NP, the 2.3.4.4b viruses contained a serine (S), while the other viruses contained an asparagine (N). Position 450 is considered variable among IAV strains, avian isolates typically contain a S or N, whereas human/mammalian strains contain a glycine (G) [39, 40]. Unexpectedly, there were no differences in the PB2 protein. Currently, the PB2 residues that interact with Mx proteins are unknown. Three residues in PB1 (59, 375, and 694) and one residue in PA (400) were also variable. Positions 375 and 694 in PB1 have been implicated as sites for avian to human adaptations, while PA 400 is thought to play a role in promoter binding [41–43]. It is possible one or more of these residues interact with Mu Mx1, which could interfere with its antiviral ability. Mammalian Mx proteins act as a barrier for avian to mammalian transmission and understanding the residues involved is important for AIV control.

Recent advancements in genome editing technology have spearheaded the development of transgenic chickens with the hope of combating AIV infection. Several strategies have been employed to create AIV resistant chickens including, using short-hairpin RNAs to interfere with AIV replication, introducing a specific gene edit to impede ANP32A function, and introducing an innate antiviral signaling transgene (RIG-I) to enhance the innate immune response [44–46]. The success of each of these strategies was variable with chickens succumbing to disease, breakthrough infections at high inoculum doses, and/or AIV mutations to overcome host boundaries [44–46]. However, many targets for enhancing innate immunity have yet to be examined. Chickens have fewer viral specific toll-like receptors (TLR) than humans or mice, a possible strategy undergoing investigation is to introduce additional TLRs into the chicken genome to enhance antiviral sensing [47, 48]. With the ability to target the virus, the host, or add a transgene there are infinite possibilities to investigate.

## Conclusion

This work proposes the use of Mu Mx1 in chickens to improve disease resistance to AIV. While Mu Mx1 was successful in reducing titers *in vitro*, *in vivo* studies are needed to confirm its effectiveness. It is also very likely that multiple strategies will be required to make a true disease-resistant bird. Gene edited poultry may be the most effective way to combat AIV when vaccination and biosecurity measures fail, as disease-resistant chickens would have significant impact on animal and human health.

## Acknowledgements

The authors acknowledge the excellent technical assistance of Mr. Ryan Sweeney. The pMAT-miniTol2 plasmid was a kind gift from Dr. Tim Doran at CSIRO in Australia. We gratefully acknowledge all data contributors for generating the genetic sequence and sharing via the GISAID Initiative.

## Author contributions

**Kelsey Briggs** Conceptualization, Methodology, Formal analysis, Writing – original draft, preparation, Writing – review & editing. **Klaudia Chrzastek:** Conceptualization, Methodology, Formal analysis. **Karen Segovia:** Conceptualization, Methodology, Formal analysis. **Jongsuk Mo:** Conceptualization, Methodology, Formal analysis. **Darrell R. Kapczynski** Conceptualization, Methodology, Formal analysis, Writing – original draft, preparation, Writing – review & editing.

## Data availability

The data are available from the senior author upon request.

