## Supplemental for "Genetic insertion of mouse Myxovirus-resistance gene 1 increases innate resistance against both high and low pathogenic avian influenza virus by significantly decreasing replication in chicken DF1 cell line"

### Supplemental Figures

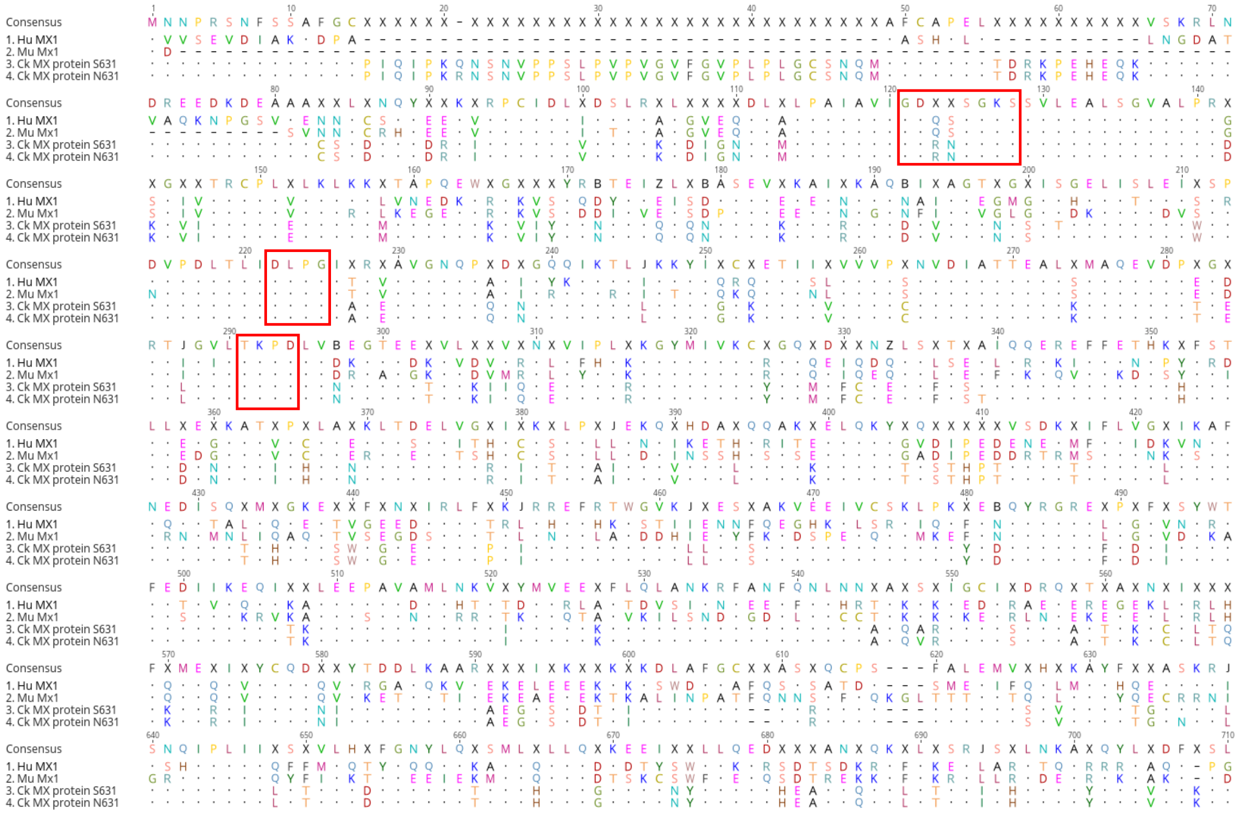


**Supplemental Figure 1: Alignment of Mx proteins.** Human (Hu), Mouse (Mu), and Chicken (Ck) Mx protein sequences were obtained from Genbank and aligned using a MUSCLE alignment in Geneious Prime. A Ck Mx protein containing S631 and N631 were aligned to compare sequences. Red boxes denote the GTP-binding consensus motifs. A dot represents a conserved residue.

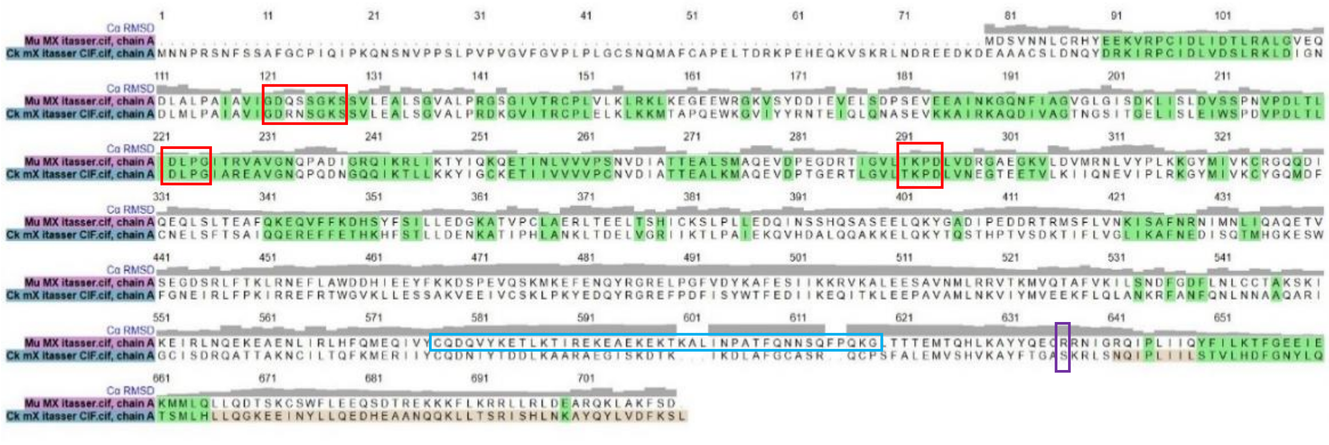


**Supplemental Figure 2. Alignment used to overlay Mu Mx1 and Ck Mx in ChimeraX for Figure 7.** Sequences in green have homologous structure and were labeled in black in Figure 7. The red boxes denote the GTP-binding consensus motifs. The light blue box denotes the L4 loop in Mu Mx1. The purple box denotes residue 631 in Ck Mx.

| | Hu MX1 | Mu Mx1 | Ck MX protein S631 | Ck MX protein N631 |
|---|---|---|---|---|
| Hu MX1 | | 244 | 393 | 393 |
| Mu Mx1 | 63.31% | | 412 | 412 |
| Ck MX protein S631 | 44.41% | 41.97% | | 6 |
| Ck MX protein N631 | 44.41% | 41.97% | 99.15% | |

**Supplemental Table 1. Number of differences and percent identity of Mx proteins.** The number of differences and percent identity was calculated based on the MUSCLE alignment. The top values correspond to the number of differences between the proteins and the bottom values are the percent identity.
